# Quantification of very low-abundant proteins in bacteria using the HaloTag and epi-fluorescence microscopy

**DOI:** 10.1101/237248

**Authors:** Alessia Lepore, Hannah Taylor, Dirk Landgraf, Burak Okumus, Sebastián Jaramillo-Riveri, Lorna McLaren, Somenath Bakshi, Johan Paulsson, M. El Karoui

## Abstract

Cell biology is increasingly dependent on quantitative methods resulting in the need for microscopic labelling technologies that are highly sensitive and specific. Whilst the use of fluorescent proteins has led to major advances, they also suffer from their relatively low brightness and photo-stability, making the detection of very low abundance proteins using fluorescent protein-based methods challenging. Here, we characterize the use of the self-labelling protein tag called HaloTag, in conjunction with an organic fluorescent dye, to label and accurately count endogenous proteins present in very low numbers (<7) in individual *Escherichia coli* cells. This procedure can be used to detect single molecules in fixed cells with conventional epifluorescence illumination and a standard microscope. We show that the detection efficiency of proteins labelled with the HaloTag is ≥80%, which is on par or better than previous techniques. Therefore, this method offers a simple and attractive alternative to current procedures to detect low abundance molecules.

## INTRODUCTION

Cell biology increasingly relies on quantitative microscopic labelling methods that provide strong fluorescent signals but are also well vetted in terms of detection efficiency and specificity. While fluorescent proteins (FPs) such as the green fluorescent protein (GFP) have become standard, they also have several disadvantages, including their relatively low brightness and photo-stability, plus the tendency to promote protein oligomerization^1^. An attractive alternative is the use of self-labelling protein tags, such as HaloTag^2^ or SNAP tag^3^, which covalently bind organic dyes. Such self-labelling tags have been widely used in eukaryotic cells for visualization in different cellular and sub-cellular compartments^4,5^. More recently, these self-labelling tags have been adapted for use in live and fixed bacterial cells for visualization^6^ and super-resolution studies^7^. Additionally, the superior brightness and photo-stability of the organic dyes make them excellent tools for super-resolution and single-particle tracking studies^8^. However, the use of such dyes in live bacterial cells is often limited by the low permeability of the bacterial envelope to the dye-substrate conjugate. This can result in low labelling efficiency of the construct, potentially limiting the use of this technology as a quantitative tool because only a partial subset of the tagged molecules is then detectable. This raises the challenge, as noted previously^7^, of whether the use of self-labelling tags can be optimized for quantitative measurements of proteins levels in bacteria.

Of particular interest is the quantification of low abundance proteins which are often important regulatory proteins^9^. Depending upon growth conditions, at least 10% of the *Escherichia coli* proteome consist of proteins that are present in less than 10 copies per cell^10,11^. This makes detection using FP fusions extremely challenging as the specific signal is barely detectable above cellular auto-fluorescence^12^. Many of these low abundance proteins have key regulatory roles, and fluctuations in their levels can impact their direct activity and all downstream processes. Therefore, it is important to accurately quantify these molecules. Ideally, for such low abundance proteins, direct counting of individual molecules is necessary as quantification of total fluorescence can introduce significant artefacts^13^. However, cytoplasmic proteins diffuse quickly within the bacterial cell and this precludes single-molecule visualization and counting by conventional epi-fluorescence microscopy. Several strategies have been devised to overcome this problem, either by using ultra-fast (stroboscopic) illumination^14^ or by reducing the diffusion of the protein of interest by localizing it to the cell membrane^15^. Other strategies are based on the chemical fixation of cells but even the gentlest procedure permanently degrades some fluorescent signal and introduces measurement noise^11^. We recently developed a method called Microfluidics Assisted Cell Screening (MACS) in which mechanical pressure is applied through a microfluidic device, significantly reducing the diffusion of cytoplasmic proteins and, thus, facilitating detection^13^. Whilst this method allows for automated microscopy at high throughput, it necessitates a specific microfluidic set-up and requires additional equipment (e.g. pressurized valves) to apply pressure on cells. Moreover, all of the above-mentioned methods need to be combined with a laser-based microscopy set-up to detect single FPs which can be expensive and is not widely available.

In this study, we combined translational fusions to the self-labelling HaloTag with chemical fixation of the cells to provide an easy-to-use quantitative tool for the quantification of protein levels across a wide expression range. In particular, we show that we can count individual very low copy proteins (<7) with very high accuracy. The HaloTag is a 33 kDa modified haloalkane dehalogenase which has been designed to covalently bind synthetic ligands comprised of a chloroalkane linker attached to a range of organic fluorescent dyes or other moieties^13^. The HaloTag ligand conjugated to the TMR dye has been described to be cell permeable^6^, thus allowing protein labelling in live cells. We show that we can use the self-labelling HaloTag for quantification of cytoplasmic protein levels in *E. coli* and it can be used to count low abundance proteins with high accuracy. Because TMR is an organic fluorophore and resists chemical fixation, we take advantage of its brightness and photo-stability to detect single molecules using a standard epifluorescence microscope set-up with a conventional LED system for fluorescence excitation. This method offers an attractive alternative to current techniques for detection and counting of proteins because it does not require a specialized microscopy set-up.

## RESULTS

### Specific *in vivo* detection of cytoplasmic HaloTag using a fluorescent ligand

We first determined conditions for *in vivo* labelling in the HaloTag in *E. coli* using the commercially available TMR ligand (Halo-Tag-Ligand TMR, here referred to as HTL-TMR)^2,16^. Bacterial strains containing a cytoplasmic HaloTag protein expressed from a medium copy-number plasmid under the control of an arabinose inducible promoter were grown in a low auto-fluorescence medium to mid-exponential phase (between OD_600_ 0.2 – 0.3). HaloTag expression was induced by adding 1% arabinose for 1 hour, HTL-TMR was subsequently added at 5μM final concentration and the cells were incubated for an additional one hour at 37°C. We chose this concentration of HTL-TMR (which is 5 times higher than concentration recommended by manufacturer^16^) to ensure complete labelling of all HaloTag proteins. After five washes of the cells with fresh growth medium to eliminate the non-bound TMR and reduce fluorescent background due to the excess of the dye, the cells were imaged by epifluorescence microscopy (Fig. 1A and Material & Methods). We observed specific labelling of the cells that contained the HaloTag protein and incubated with HTL-TMR. Virtually no signal was detected in normal wild-type cells (i.e. cells that contained no HaloTag and incubated with HTL-TMR) or in cells carrying the HaloTag protein that were not incubated with HTL-TMR (Fig. 1B). These results indicate that specific detection of cytoplasmic proteins can be achieved using *in vivo* labelling of the HaloTag with HTL-TMR, thus confirming previous reports^6^.

**Figure 1.**
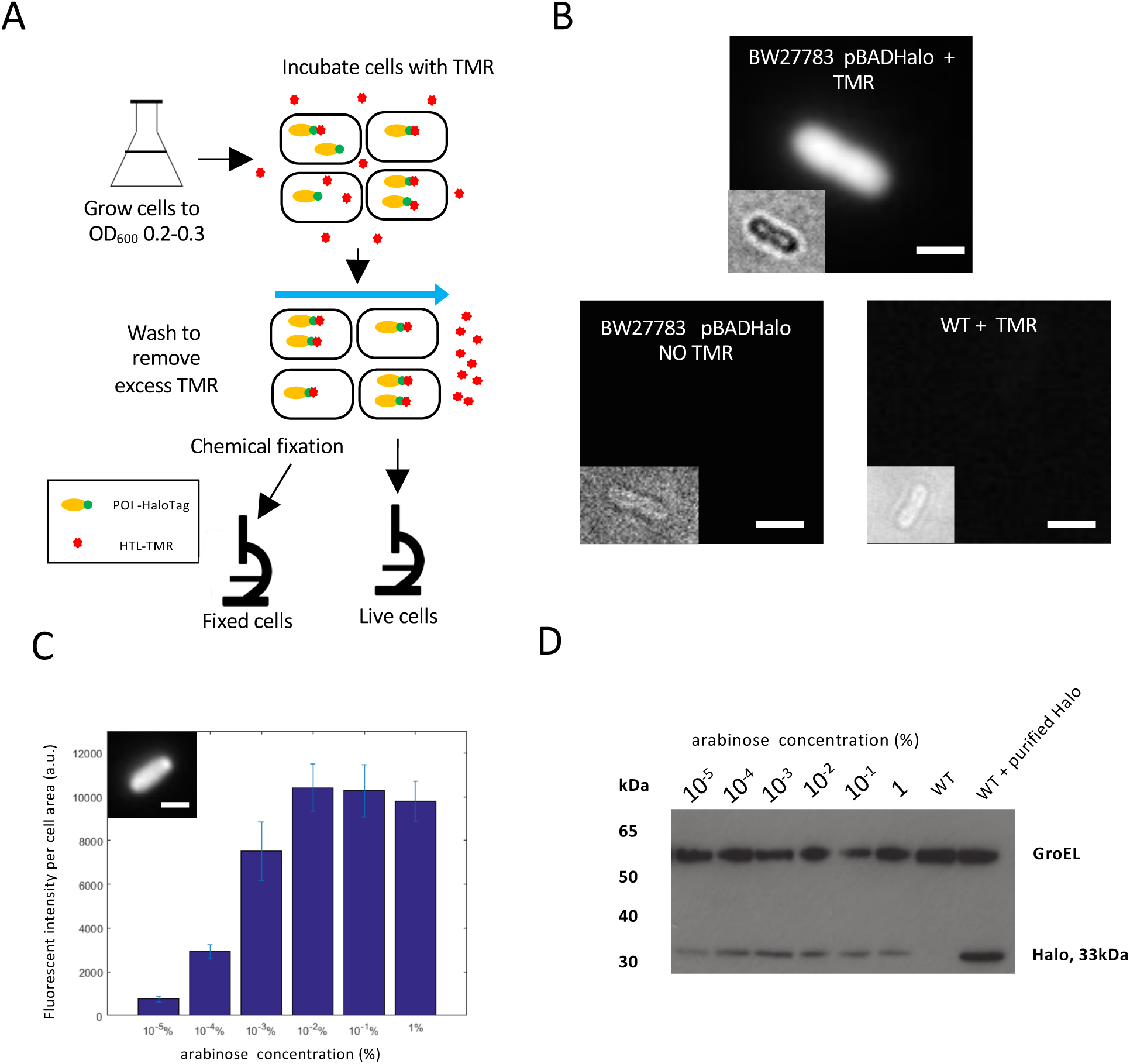
Labelling of the HaloTag is specific and allows quantitative detection of protein levels. A. Outline of the protocol used in this study. Bacterial cells were grown overnight, freshly diluted and grown until an O.D_600_ of approximately 0.2. They were then incubated in HTL-TMR for one hour followed by several washes. Live cells were then either immediately visualized by microscopy or chemically fixed before microscopy. B. Specific detection of labelled HaloTag molecules in live cells. A strain expressing the HaloTag protein (arabinose concentration 1%) was incubated with HTL-TMR and showed an intense fluorescent signal (top). Virtually no signal was detected in a strain expressing the HaloTag protein not incubated with HTL-TMR (bottom left) and or in a wild-type strain incubated with HTL-TMR (bottom right). All the images are displayed using the same minimum and maximum intensity values. Insets: corresponding bright-field images defocused to identify cell contour. Bar = 1 micron. C. The HaloTag protein expression from an arabinose inducible promoter is proportional to the arabinose concentration. The fluorescence per cell area was plotted as a function of arabinose concentration. Cells were chemically fixed immediately after washing to stop further expression of the HaloTag protein. Data correspond to the mean of at least two experiments and error bars correspond to the standard deviation of the mean. Inset: example of fixed cell expressing Halo using 1% arabinose in the growth medium. Bar 1μm. D. Western blot detection of HaloTag expression in the same induction conditions. Total cell lysates where prepared and probed with an antibody specific for HaloTag (Promega). This analysis showed increased HaloTag expression with increased arabinose concentration in good agreement with the quantification of the microscopy data. The GroEL is shown below to indicate that each lane has been loaded with a similar amount of protein. The full-length Western blot is presented in Supplementary Figure 14.

### Labelling of the HaloTag with TMR enables measurements of protein levels

To test whether Halo-TMR labelling could be used to quantify variation in protein levels, we measured the average fluorescence intensity of cells expressing the HaloTag protein in the presence of various arabinose concentrations. After *in vivo* labelling of the protein and washing of the cells, we chemically fixed the cells to stop further protein production which would lead to under-estimation of protein concentration as newly synthetized proteins would not be labelled. As expected, we observed that fluorescence intensity per area increased with increasing concentration of arabinose, indicating that Halo-TMR labelling correctly reports increased protein production (Fig. 1C). Quantification of the average fluorescence intensity indicated that when the inducer concentration varied from 0.00001% to 0.1%, the average fluorescent intensity of the population increased by approximately 10 fold, similar to levels previously observed for a construct where GFP was driven by the same promoter on a similar plasmid^17^. At higher induction levels (1% arabinose), the fluorescence intensity decreased, again similar to observations using GFP^17^. Indeed, analysis of protein production by Western blot showed lower level of HaloTag protein production at 1% arabinose than at 0.1% (Fig. 1D). Analysis of the fluorescence intensity per cell showed that HaloTag expression was homogeneously distributed across the population at all concentration of inducer (Fig. S1). This is expected because we used a background strain where the positive feedback loop controlling the arabinose sensitive promoter has been disrupted, leading to homogenous expression from this promoter^17^. Taken together, these results indicate that the HaloTag labelling can be used to quantify protein levels in single cells with the same sensitivity as FP-based methods.

### Detection of an endogenous low-abundance protein

We reasoned that the brightness of the cell-permeable TMR dye in combination with the self-labelling HaloTag protein would enable the detection of proteins present in very low numbers in *E. coli* cells. To test the feasibility of this approach, we constructed a translational fusion of the HaloTag to RecB, a cytoplasmic protein involved in DNA repair that has been reported to be present in 10 or fewer copies per cell^11,18^. RecB is part of a heterotrimeric complex called RecBCD and examination of the RecBCD structure^19^ indicated that a fusion inserted in a loop situated after Ser47 would most likely not interfere with complex formation. Therefore, we introduced the HaloTag surrounded by two short linkers at this position using markerless homologous recombination^20^ to create a scarless fusion, which was introduced at the endogenous chromosomal locus of the *recB* gene and replaced the wild-type gene. We measured growth curves to compare the strain harbouring the RecB-HaloTag fusion to the wild-type strain, and the data indicated that the tagged strain was not impaired for growth (Fig. S2). We confirmed that the RecB-HaloTag fusion was fully functional for DNA repair by showing that the tagged strain was as resistant as the wild-type to nalidixic acid, a DNA damaging agent that leads to at least 10^3^ fold reduction in viability when RecBCD is inactive^21^ (Fig. S3). To test the integrity of the fusion protein, a Western blot of total cells lysates was performed and probed with anti-HaloTag antibody (Fig. S4). We observed a single band of the expected size for the full-length fusion, indicating that there was no proteolytic processing of the fusion protein.

Given that RecB is expressed at very low levels, epifluorescence imaging of live cells with the TMR-labelled HaloTag leads to very weak diffuse signal that is difficult to detect above cellular auto-fluorescence background. However, as HTL-TMR is resistant to chemical fixation, we reasoned that we could take advantage of its brightness and photo-stability to detect single molecules of RecB-Halo after chemical fixation. As molecules do not move in fixed cells (Fig S5), long exposure times can be used in epifluorescence microscopy to allow detection of a distinct diffraction-limited fluorescent spot for each individual molecule. Therefore, we detected endogenous RecB molecules in cells that were chemically fixed after *in vivo* labelling with HTL-TMR. Since RecB does not homo-oligomerize, individual RecB-HaloTag fusions are labelled with only one HTL-TMR dye per protein molecule. As shown in Fig. 2A, when a 1-second exposure time was used, 5-10 diffraction-limited spots per cell were detected, where each fluorescent spot corresponds to an individual RecB-HaloTag molecule. Indeed, analysis of the fluorescence bleaching patterns of the spots confirmed that each spot corresponds to a single molecule as we observed abrupt disappearance of fluorescence in a single step (Fig. 2B). Thus, we concluded that TMR labelling of cytoplasmic proteins tagged with the HaloTag allows detection of single molecules in fixed cells using conventional epifluorescence microscopy.

**Figure 2.**
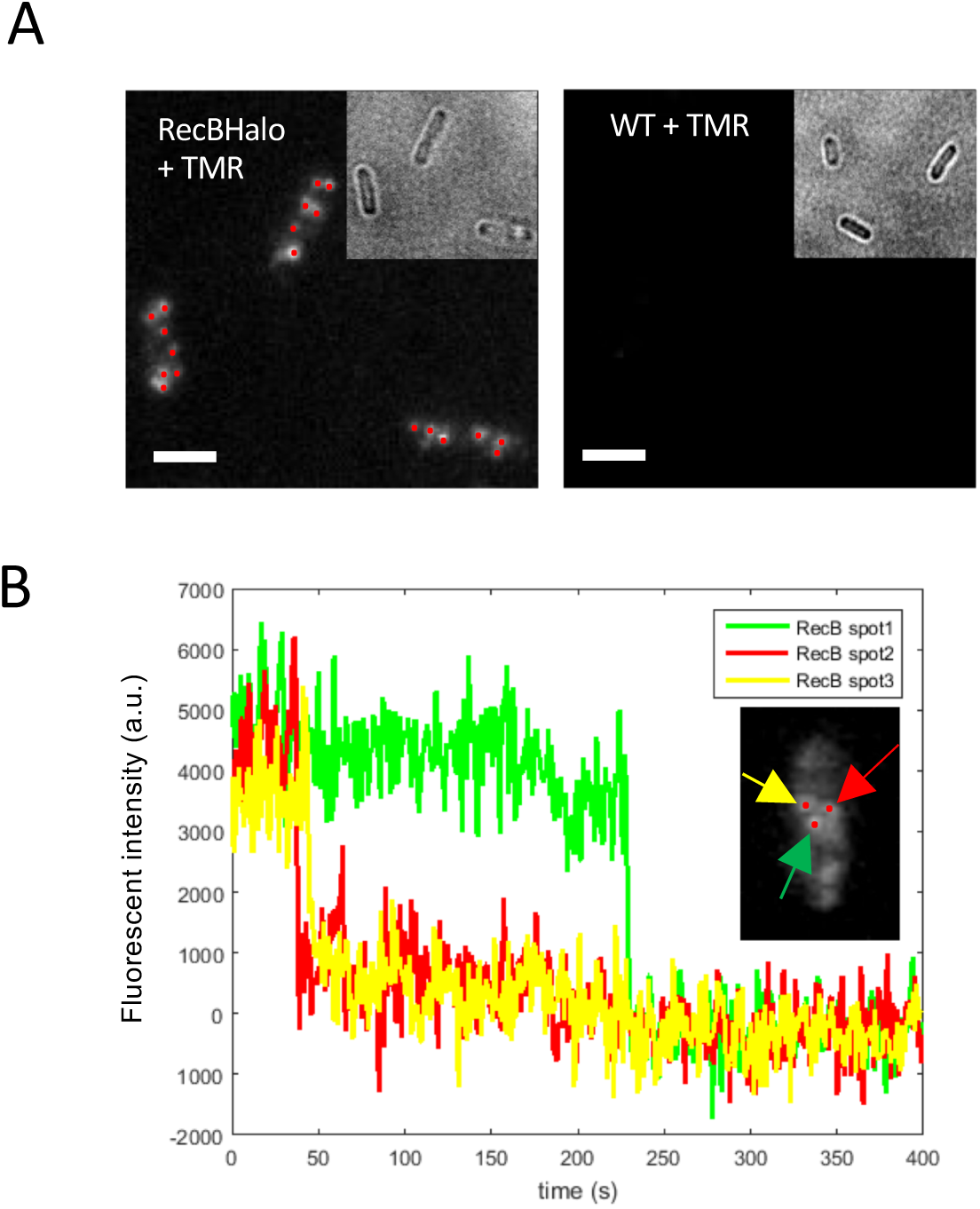
Single molecule detection of a low abundance protein. A. HaloTag-HTL-TMR allows single molecule detection. After *in vivo* labelling cells were chemically fixed to stop protein diffusion. A strain containing a RecB-HaloTag fusion showed diffraction limited spots corresponding to single RecB molecules when incubated with HTL-TMR (left panel). The WT strain incubated with TMR did not shows any signal (right). The maximum intensity projection of the z-stack images (from the 2^nd^ to the last image) is shown. All the images are displayed using the same minimum and maximum intensity values. B. Single molecule detection. Representative time traces of RecB-HaloTag spots from same cell (after background subtraction) exhibited single-step photobleaching indicating detection of a single fluorophore.

### Counting of RecB molecules using the HaloTag fusion

As cytoplasmic molecules stop moving after chemical fixation, the resulting fluorescent spots can easily be counted, thus providing a method for single molecule counting of low abundance proteins. We quantified the number of RecB-HaloTag molecules using a spot detection algorithm that detects the diffraction-limited spots within a single cell and counts them (see Material and Methods, and Fig. S12). The number of molecules of a particular protein is expected to double during the cell cycle as cells about to divide have on average twice more proteins that newborn ones. To avoid bias when estimating of the number of molecules per cell, we quantified spots only in newborn cells by conditioning on cell size. We observed that newborn cells had on average 4.9±0.3 RecB-HaloTag molecules per cell (Fig. 3A). In contrast, we only detected 0.2 spots on average per cell in a strain that did not express the HaloTag, which further confirmed the specificity of our labelling method. We note that we detected more RecB molecules per cell than previously reported in a large-scale quantification of the *E. coli* proteome, which reported 0.6 RecB molecules per cells^11^. This is possibly because that study used a C-terminal fusion of Venus-YFP to RecB. Such a fusion is very likely to disrupt RecBCD complex formation which could result in a phenotype similar to a *recBCD* mutant. These mutants have very low viability^22^, which would lead to under-estimation of the average number of molecules per cell as many cells are actually unable to grow ^23^. However, the number of RecB molecules we observed (from 2 to 9 per cell) is in good agreement with the number of 9.5 reported in a recent population-based high-resolution mass spectrometry quantification^24^.

**Figure 3.**
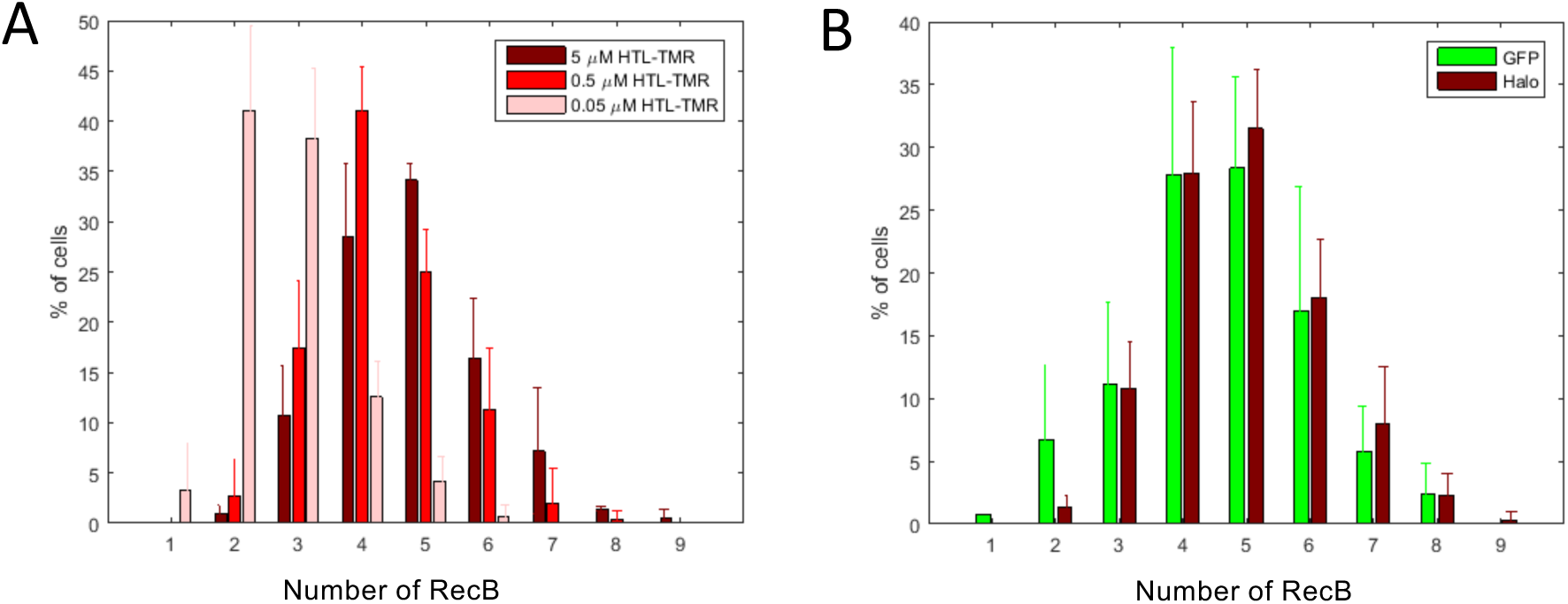
Single-molecule counting of RecB-HaloTag. A. HTL-TMR concentration impacts the labelling efficiency. Absolute numbers of detected RecB-Halo molecules per cell for three HTL-TMR concentrations are depicted. The numbers were measured for newborn cells only. When incubated with low concentration of HTL-TMR (0.05μM) only a subset of all RecB-Halo molecules was detected. The proportion of cells depicted are the mean of three experiments (total number of cells 261, 226 and 231 for 5μM, 0.5μM and 0.05μM respectively) and the error bar correspond to the standard deviation of the mean. B. Comparison of RecB number distribution using Halo-Tag based detection or MACS based detection with the RecB-sfGFP fusion. RecB spots were detected either with RecB-HaloTag incubated with 5μM HTL-TMR and chemical fixation or with a RecB-sfGFP fusion detected by HILO microscopy combined with mechanical slowing down of molecules using MACS^13^. The proportion of cells depicted are the mean of three and five experiments for RecB-sfGPF and RecB-HaloTag, respectively (total number of cells 109 and 241) and the error bar correspond to the standard deviation of the mean.

To further validate our labelling, we then tested the dependence of the detection efficiency on the concentration of HTL-TMR dye used for the labelling. We incubated the strain carrying the RecB-HaloTag fusion with final concentrations of HTL-TMR ranging from 0.05μM to 5μM for an hour before washing and fixing the cells. Quantification of the detected fluorescent spots indicated that a lower number of spots (on average 2.9±0.3) were detected when the cells were incubated with 0.05μM HTL-TMR, suggesting that under these conditions, the dye is limiting for the labelling reaction (Fig. 3A). In contrast, we detected almost identical average number of spots when the cells were incubated with 0.5μM or 5μM HTL-TMR indicating that these concentrations were sufficient for accurate detection. To ensure maximum detection, we decided to use the higher concentration of HTL-TMR (i.e. 5μM) since it could be used without increasing false positive detection.

### Evaluation of the detection efficiency of the Halo tag-based method for low abundant protein

To evaluate the detection efficiency of this labelling method, we compared HaloTag-based detection with our recently developed MACS method, which uses a microfluidic device for mechanical fixation of freely diffusing cytoplasmic proteins and, therefore, enables the detection of single fluorescent proteins in live cells^13^. We built a fusion of RecB to sfGFP (sfGFP inserted after Ser47 in the same position as HaloTag) and confirmed that the fusion was functional. We compared the number of RecB-HaloTag molecules detected by both methods and observed an excellent agreement (Fig. 3B). The average copy number of RecB molecules per cell was 4.5 ± 0.4 for MACS with the RecB-sfGFP fusion and 4.9 ± 0.3 with RecB-HaloTag fusion. A two-sample Kolmogorov-Smirnov test performed on the distributions of RecB-Halo and RecB-sfGPF showed that they are similar within a significance level of 5% (Fig. S6). We observed a shift towards slightly higher values for the HaloTag based method, indicating that, at least when using the high HTL-TMR concentration, this method may be slightly more sensitive. This is possibly a result of limitations of FP-based methods because of the maturation time of the protein. We previously determined that the MACS based method detects at least 80% of the tagged proteins^13^, which suggests that the detection efficiency of the HaloTag based method is 80% or greater. To further evaluate the detection efficiency of this method, we built a fusion of RecB with two Halo tags positioned in tandem and separated by a 31 amino-acid linker. As for others constructs, we checked that the fusion did not impair RecB and strain viability (Fig S2, S3). If labelling with a single Halo-tag resulted in a low level of detection, one would expect a significantly higher number of spots when the protein is labelled with two Halo-tags. In contrast, we observed similar distributions when detecting RecB with a single or a double Halo-tag (on average 5.0±1.2 and 4.6±1.3 molecules were detected, Fig. S7). The similarity of the distributions was confirmed by a two sample K-S test within a significant level of 5% (Fig. S8).

To test the validity of this method on another protein, we built a fusion of RecC to HaloTag. RecC is part of the same complex as RecB but is transcribed from another promoter and has been reported to be expressed at a similar level to RecB ^25,26^. We checked that the fusion of RecC to HaloTag did not compromise the functionality of the protein (Fig S2, S3 and S4). We compared the HaloTag-based quantification of RecC with RecC-sfGFP quantification performed with MACS^13^. We observed that the average number of molecules per cell (for cells of length < 3.5 μm) was 4.6 ± 1.1 RecC-sfGFP using MACS, and 4.7 ± 1.1 for RecC-HaloTag (Fig S9). The similarity of the distribution was confirmed by a two-sample Kolmogorov-Smirnov test (Fig S10).

Thus, we conclude that our new method offers a versatile, easy and highly quantitative alternative to FP-based single molecule counting techniques.

## Discussion

In this work, we demonstrated the use of the self-labelling HaloTag for quantification of protein levels in bacterial cells over a wide range of expression levels. In particular, we show that for low-abundance proteins, we can detect more than 80% of molecules showing similar or better detection capabilities than FP-based methods. The HaloTag-based procedure is simpler than most single molecule counting methods because the cells can be chemically fixed and imaged with a conventional epifluorescence microscope.

However, there are some potential limitations to this method. Firstly, precise detection of individual molecules is limited to low abundant proteins: due to the small size of bacteria, even for a relatively small number of particles the inter-particle distance can be quite small and often below the diffraction-limit leading to undercounting artifacts. We evaluated this effect using simulations (see Materials and Methods and Fig. S13) and showed that undercounting has a very limited effect on the average number of molecules if there are less than 6.3 proteins per cell. For more abundant proteins, detection of single step photobleaching can be used to identify spots that correspond to more than one molecule ^27^. Preliminary observations suggest this could be used in the future to extend the range of our method, as when labelling the RecB-two Halo tags fusion we observed photobleaching curves with two steps, consistent with two fluorophores binding the two Halo protein fusion. Contrary to FP-based methods where cells can be immediately visualized, our method requires first labelling and then repeated washes to remove the unbound dye molecules. Proteins produced after the labelling will not be detectable which may lead to under-estimation of the total number of molecules present, although this limitation can be alleviated by chemically fixing the cells immediately after labelling.

Secondly, we have shown that high concentrations of HTL-TMR are necessary to achieve a high labelling efficiency and it is likely that the ideal labelling concentrations must be adjusted for different bacterial species (because of the differences of membrane permeability properties or the presence of efflux pumps that may reduce the intracellular concentration of HTL-TMR). However, this limitation can also be used as an advantage: we show that by using lower dye concentration, it is possible to obtain “under-labelling conditions” where only 2– 3 molecules are labelled on average, which facilitates single molecule tracking in live cells. In fact, the use of HaloTag-based labelling for *in vivo* single-molecule tracking has been recently demonstrated in *Salmonella enterica*^7^ and these types of applications are likely to expand further in the future. Indeed, the development of new HaloTag compatible fluorophores (such as JF549) that are even brighter and more photo-stable, plus photo-activatable dyes, will greatly facilitate the use of self-labelling tags for single molecule tracking or super-resolution applications^7,8,28^. As a proof of principle, we have confirmed that the new Janelia Fluor 549 dye (JF549) is suitable for live labelling in *E. coli*. We labelled a RecB-HaloTag strain *in vivo* with JF549 using the same protocol that we developed for HTL-TMR. We then counted the number of detected molecules after fixation and obtained similar detection efficiency than with HTL-TMR. Moreover, we noted that this newly developed dye leads to fluorescent spots that are easier to detect, thus facilitating counting (Fig. S11).

In conclusion, we have developed a method that is suitable for counting low abundance proteins in bacterial cells in a fast and reliable manner. Our method provides a very attractive alternative to FP-based labelling methods and it can be used as an independent validation method to confirm results obtained with FP protein fusions. In addition to its immediate application as a counting method, the quantitative information provided is directly relevant for single molecule tracking experiments and is compatible with new dyes that are developed for such applications.

## Materials and Methods

### Culture conditions

For all microscopy based experiments, the cell cultures were grown in M9 supplemented by 10% LB, with final concentration of 1X M9 salts, 10% (v/v) LB, 0.2% (w/v) glucose, 2 mM MgsO_4_ and 0.1 mM CaCl_2_. We refer to this medium as “imaging medium”. When necessary, chloramphenicol was used at a final concentration of 30µg/ml. In induction experiments, the medium was supplemented with various concentration of arabinose. For Western blot experiments, cells were grown in LB.

### Strain and plasmid construction

*E. coli* MG1655 was used as WT strain in this study except for the experiment using Halo under the control of the arabinose promoter which used BW27783 as a background strain. The characteristics of the strains used in this study are described in Table 1. Oligonucleotide sequences are available in Supplementary Table 1.

**Table 1.** Strains and plasmids used.

| Name | Characteristics | Reference |
| --- | --- | --- |
| Bacterial strains |  |  |
| MG1655 | *F-lambda-rph-1* | Lab stock |
| BW27783 | $\Delta(araFGH) \phi(\Delta araEpPCP8-araE)$ | 17 |
| MEK65 | MG1655 *recB::halotag* | This work |
| MEK705 | MG1655 *recB::sfGFP* | This work |
| MEK573 | MG1655 *recC::halotag* | This work |
| MEK711 | MG1655 *recC::sfGFP* | This work |
| MEK556 | MG1655 *recB::halo::cohalo* | This work |
| Plasmids |  |  |
| pSF1 | Expression of HaloTag under the control of the pBAD promoter | This work |
| pBH35 | Template for plasmid mediated recombination to construct *recB::halotag* fusion | This work |
| pBH29 | Template for plasmid mediated recombination to construct *recB::sfGFP* fusion | This work |
| pHT8 | Template for plasmid mediated recombination to construct *recB::halo::cohalo* fusion | This work |

Plasmid pSF1 was constructed to allow expression of the HaloTag under the control of an arabinose inducible promoter. pBAD33^29^ was digested using SacI and PstI restriction enzymes. An insert containing a Ribosome Binding Site, the HaloTag open reading frame and a stop codon was amplified from pBH36 (this work) using primers OSF1 and OSF2. The digested PCR product was inserted into the cut vector using isothermal assembly^30^ and the mix transformed into *E. coli* DH5α. Successful plasmid construction was checked by colony PCR and sequencing.

Plasmid pBH35 was created in order to build an in-frame fusion of *recB* to *haloTag*, with *haloTag* (separated from *recB* by two short linkers) being inserted after Ser47 of *recB*, thus creating strain MEK65. This plasmid enables direct replacement of the wild-type gene with the fusion at the endogenous chromosomal locus by plasmid mediated recombination^20^. In brief, the pTOF24 vector was linearized by SalI/PstI digestion and three PCR fragments were assembled by isothermal assembly into this vector^30^. The fragments were respectively amplified with the oligos Obh41/Obh67 (fragment containing homology to *recB* before Ser47), Obh66/70 (fragment containing the linkers and the *halotag* gene) and Obh69/46 (fragment containing homology to *recB* after Ser47). Correct plasmid construction was verified by colony PCR and sequencing. Plasmid pBH29 was created in order to build an in-frame fusion of *recB* to *sfGFP*, with *sfGFP* (separated from *recB* by two short linkers) being inserted after Ser47 of *recB*, thus creating strain MEK705. This plasmid enables direct replacement of the wild-type gene with the fusion at the endogenous chromosomal locus by plasmid mediated recombination^20^. In brief, the pTOF24 vector was linearized by SalI/PstI digestion and three PCR fragments were assembled by isothermal assembly into this vector. The fragments were respectively amplified with the Obh41/Obh42 oligos (fragment containing homology to *recB* before Ser47), Obh43/44 (fragment containing the linkers and the *halotag* gene) and Obh45/46 (fragment containing homology to *recB* after Ser47). Correct plasmid construction was verified by colony PCR and sequencing.

Plasmid pHT8 was constructed in order to build an in-frame fusion of *recBhalo* with a second *halotag* (separated from the *recBhalo* by a linker of 31 amino acid) being inserted after the first *halotag*, thus creating MEK556. In brief, the pTOF24 vector was linearized by SalI/PstI digestion and three PCR fragments were assembled by isothermal assembly into this vector. The fragments were respectively amplified with the oligos Oht82/Oth83, Oht86/Oht87. Correct plasmid construction was verified by colony PCR and sequencing.

The construction of the *E. coli* strains with translational fusions to RecC in C-terminal at its endogenous locus was performed with the lambda red method^31^, as previously described^1^. Primers with 40-nucleotide upstream or downstream homology with *recC* (0bh35 and Obh36) were used to PCR amplify the respective integration cassettes using pDHL502 (sfGFP), pDHL940 (HaloTag) as a template. Correct strain construction was verified by colony PCR and sequencing.

### *In vivo* labelling of HaloTagged proteins

Overnight cultures were inoculated in imaging medium from glycerol stock and incubated with shaking at 37°C. These cultures were diluted 1 in 250 in fresh imaging medium, incubated shaking at 37°C and the cells were grown to mid exponential phase (OD_600_=0.2 - 0.3). The equivalent of 1 mL of cells at OD_600_= 0.2 was re-suspended in fresh imaging medium supplemented with HTL-TMR or JF549 at a final concentration of 5μM (except when otherwise noted). The culture was further incubated for 1 hour at 37°C with shaking. After the labelling step, each sample was washed 4-5 times with 1 ml imaging medium. At each step, cells were transferred to a new tube to facilitate the removal of the dye. Cells were either imaged immediately or subjected to chemical fixation in which case each cell pellet was resuspended in freshly prepared fixation solution (2.5% formaldehyde, Thermo Scientific #10751395, in 1X PBS) for an hour at room temperature. The cells were pelleted and then washed with 1 ml 1X PBS twice more and mounted on an agarose pad for imaging.

### Fluorescence microscopy

Imaging was performed using an inverted microscope (Nikon Ti-E) equipped with an EMCCD Camera (iXion Ultra 897, Andor), a SpectraX Line engine (Lumencor) and a 100X Nikon objective (NA 1.45, oil immersion). RecBHalo-HTL-TMR (or JF549) was detected using a red LED and a TRITC filter cube by acquiring z-stack images (total range 0.8-1μm) of fixed cells using an exposure time of 1-2 sec. For each position, Z-stacks comprised of 4 or 5 images separated by 200nm were acquired.

### Image analysis: Spot-finding analysis

The fluorescent spots finding analysis can be summarized as a two-step procedure: finding the cells (segmentation) and counting the spots in each cell (counting).

The natural auto florescence of the cell in the green excitation channel was used to find the cells in the image. The cells segmentation procedure was based on a Matlab routine (https://uk.mathworks.com/help/images/examples/detecting-a-cell-using-image-segmentation.html). Errors in cell identification were manually corrected. Several cell characteristics such as area, perimeter, cell length and width were computed for each detected cell. To count molecules in newborn cells, we restricted the analysis to cells with a length lower than 3.5μm.

The number of diffraction-limited spots was computed using the maximum projection image from the acquired z-stacks (without the first z-stack frame that contain most of the auto-fluorescence signal). This enabled a better detection of the single spots over the background. First, an area of 30×35 pixels around the centroid of each cell was cropped. Then a band pass filter was used to remove high-frequency noise and low-frequency features. A spot was identified as a local maximum of a size of 6×6 pixels having an intensity above the local threshold. The band pass filter and peak-finder functions are from a previously developed and published software (http://physics.georgetown.edu/matlab/)^13^.

### Simulation to evaluate potential limits in the counting

The point-spread-function of the microscope sets the diffraction limit for resolving two particles (~250 nm), causing any two or more particles within this distance to appear as a single spot. There is a second source of undercounting which comes from the 2D images of the 3D spatial distribution of the particles. The particle detection algorithms use 2D projection of the 3D distribution of the particle emitters, which can cause more than one particle to appear as single spot if they are close in the image plane but separated in the optical axis.

In order to quantitatively estimate the extent of undercounting, we developed a computer simulation to assess the degree of potential undercounting. We considered the different cell sizes sampled from the distribution of cells sizes observed in the experiment. We then modelled the confinement as a spherocylinder^32^ with the chosen dimensions. N molecules (sampled from a Poisson distribution with supplied mean) were randomly placed in this virtual bacterial cell. The we used DBSCAN clustering algorithm to cluster any number of particles that are within the diffraction limit of each other. The count of spatially resolved and unresolved clusters was compared to the input N to measure the extent of undercounting.

The computer code was developed and executed in MATLAB and it is available on request.

### Quantification of total fluorescent intensity

Fluorescence images were segmented by a custom-made algorithm implemented in MATLAB. In brief, the algorithm was designed to detect pixel outliers (i.e. cells) in the distribution of intensity values. The image was background corrected by subtracting a morphologically opened version of the initial image. Next, a Gaussian filter was used to sharpen the signal and reduce the noise. All the images contained relatively sparse cells, therefore, most of the pixels did not belong to cells: an approximation of the “non-cell” intensity distribution was computed (from the low-intensity pixels) and used it to threshold the image. To further distinguish cells in close proximity, an arbitrary fraction of the lower intensity pixels was removed from each resulting connected component. Finally, segmentation results were manually curated to remove false positives or cells incorrectly segmented. Using the segmented images, the average intensity per pixel of each cell was quantified and used as a measure of protein concentration.

### Statistical Analysis

The data of the HaloTag protein expression (Fig1C) are the average of two independent experiments for each concentration and the error bars represent the standard deviation from the mean.

The distributions for the single molecule numbers are averaged over three independent experiments (except otherwise stated in the figures description) and the error bar is the standard deviation from the mean.

The two-sample Kolmogorov-Smirnov test was performed using the MATLAB function kstest2.

### Data availability

The datasets generated and analyzed during the current study are available from the corresponding author on request.

## Acknowledgements

We are grateful to Dr. J. Salje for help in analyzing RecBCD 3D structure, and to S. Fawcett and E. Jean for preliminary data. We thank Dr L. Lavis (Janelia Farm) for sharing the JF459 dye. This work has been supported by a Wellcome Trust Investigator Award 205008/Z/16/Z (to M.E.K.), a National Institutes of Health grant RO1GM081563 (to J. P.), a Darwin Trust of Edinburgh postgraduate studentship (to S.J.R), a Marie Curie Fellowship PIOF-GA-2009-254082 – DRIBAC (to M.E.K.) and a SULSA award H17014 (to A. L.).

## Author Contributions

M.E.K., H. T., D.L., and J. P. conceived the experiment. M.E.K, H.T. and McL. built the strains. H.T., A.L and M.E.K optimized the labelling protocol. M.E.K, H.T. and A.L. performed the labelling experiment. M.E.K, D.L., B.O. and A.L performed the MACS experiments. A.L. and S.J.R wrote the analysis software. L.McL. performed the Western blots. S.B. wrote and performed the simulations to assess the accuracy of the image analysis. A. L., H.T., D.L. and M.E.K. wrote the manuscript. All authors reviewed the manuscript.

## Additional Information

The authors declare no competing interests.

