## supplementary figures and tables for "Quantification of very low-abundant proteins in bacteria using the HaloTag and epi-fluorescence microscopy"

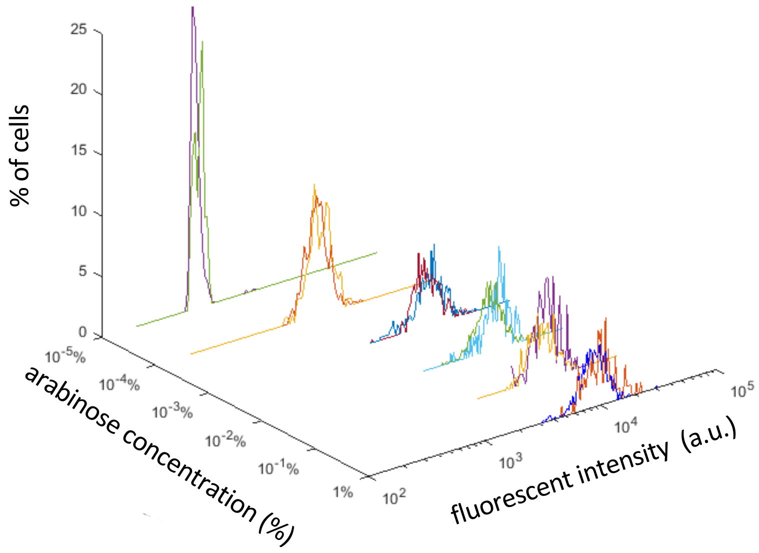


**Figure S1: Single cell expression of Halo protein after arabinose induction**. The distribution of single cell total fluorescent signal (normalized by cell area) is depicted for various arabinose concentrations. Two representative datasets for each arabinose concentration are shown.

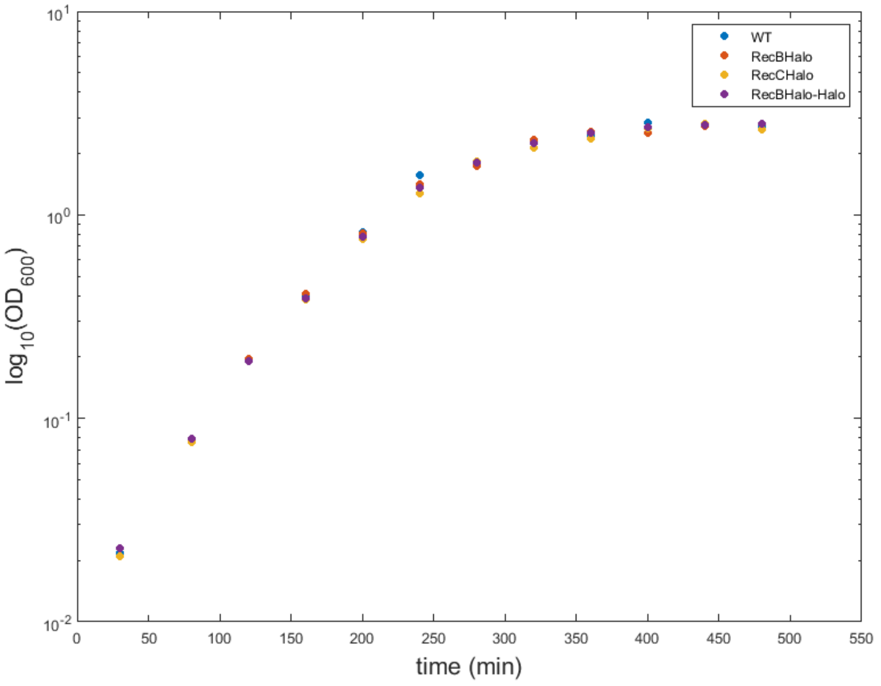


**Figure S2: All the strains built with the HaloTag fusion (RecB-Halo, RecC-Halo and RecB-Halo-Halo) have the same growth rate as the WT strain**. Growth curve of WT strain (blue), RecB-Halo fusion (orange), RecC-Halo fusion(yellow) and RecB-Halo-Halo fusion(purple). The bacteria were grown in imaging media with aeration.

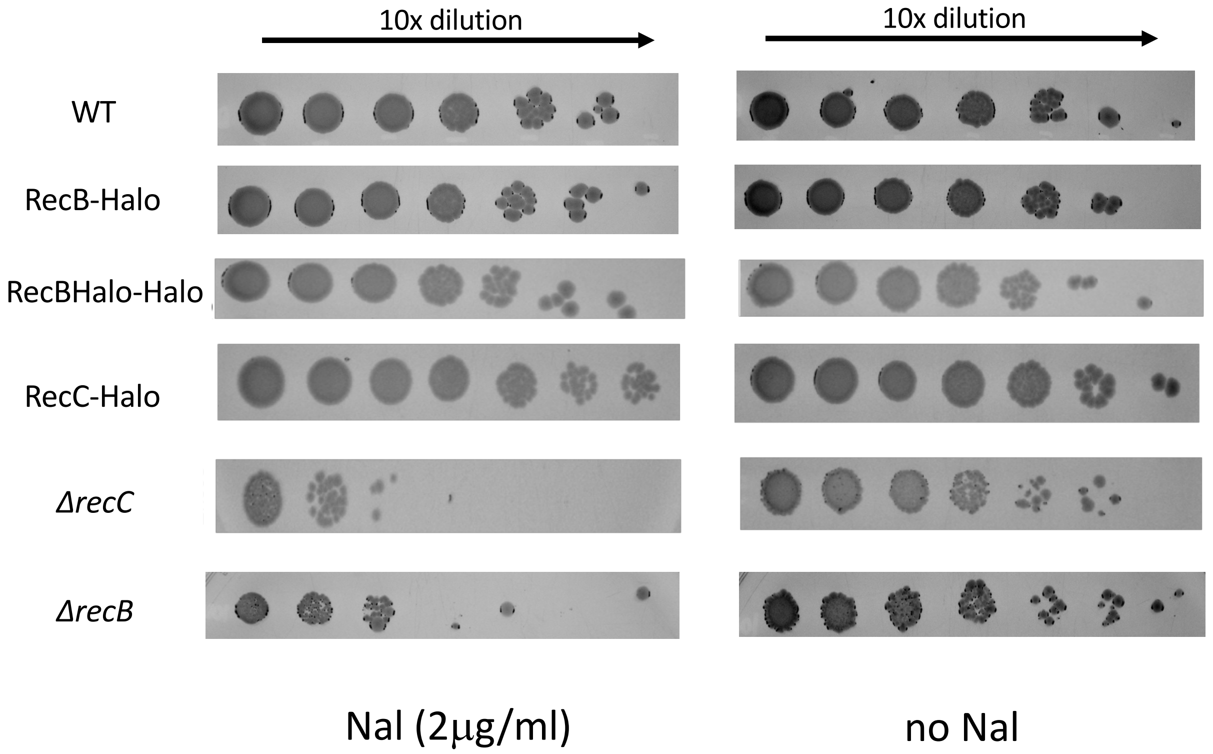


**Figure S3. The HaloTag fusions are fully functional**. Serial dilutions of WT, RecB-Halo, RecB-Halo-Halo, RecC-Halo, ΔrecC and ΔrecB strains were spotted onto LB plates containing 2$\mu$g/ml nalidixic acid or no nalidixic acid. (Nalidixic acid causes DNA damage which is repaired in a RecBCD-dependent manner). The strains carrying HaloTag fusions show the same viability as the WT whereas the ΔRecB and ΔRecC strains show a 1000 time reduction in viability compared to the WT at 2$\mu$g/ml nalidixic acid .

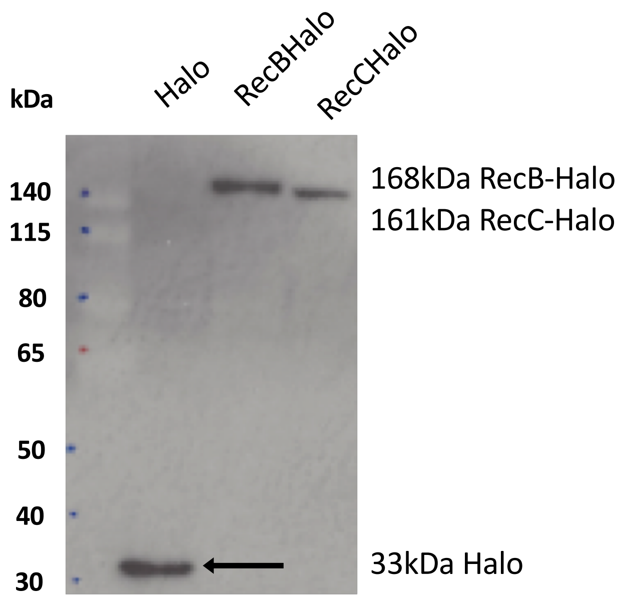


**Figure S4. RecB-Halo and RecC-Halo are expressed as a full length protein.** *E. coli* strains expressing either RecB-HaloTag, RecC-HaloTag or HaloTag only were grown in LB and protein cell lysates were analyzed by Western blot using an antibody specific to the Halo-Tag (Promega). Both the HaloTag protein, the RecB-HaloTag and the RecC-HaloTag fusion are detected by a single band at their expected size (33kDa 168 kDa and 161kDa respectively).

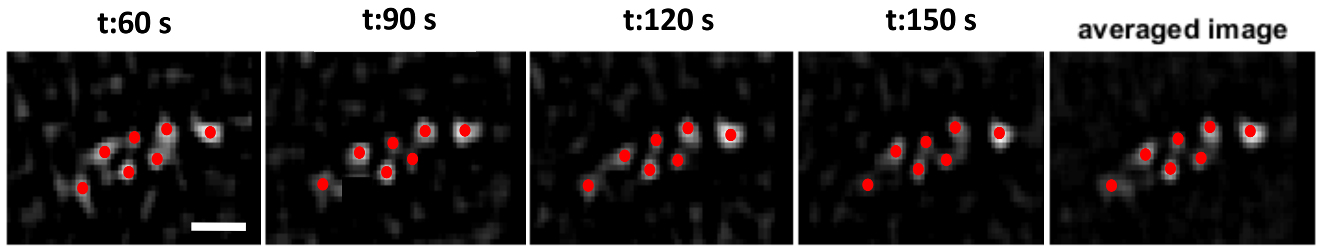


**Figure S5. Single molecule detection is not affected by chemical fixation.** Time-lapse images acquired every 30 seconds show that the number of detectable molecules and the position of the diffraction limited spots do not change over time in chemically fixed cells. The pictures presented here correspond to the filtered images of the z-stack maximum intensity projection. Each image was acquired with an exposure time of 500 ms.

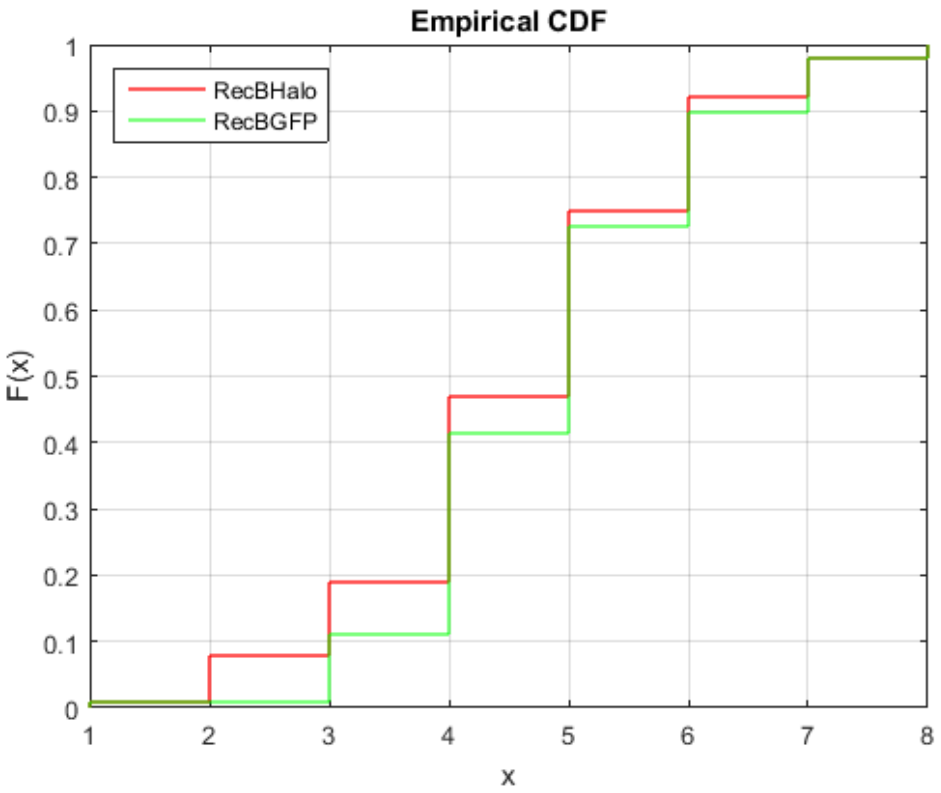


**Figure S6. RecB-HTL cumulative distribution comparisons with RecB-sfGFP.**

Empirical cumulative distribution for RecB-Halo (red) and RecB-GFP (green) are shown. A two-sample Kolmogorov-Smirnov test accepts the null hypothesis indicating that the data come from the same distribution (*p*-value = 0.90 > 0.05, the default level of significance).

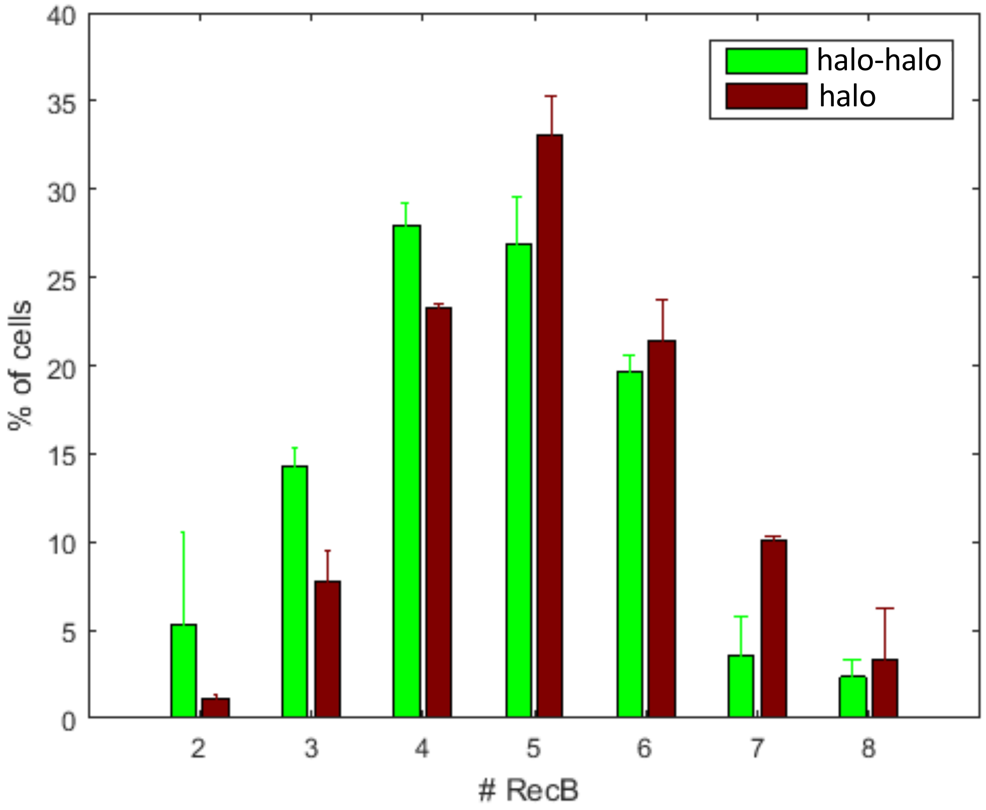


**Figure S7. Quantification of spots detected with RecB-Halo and RecB-Halo-Halo shows similar value with both fusions.**

Comparison of RecB-Halo (red) distribution and the RecB-Halo-Halo (green) using HTL-TMR based detection. Each distribution is the average of two experiments. Total number of cells for the RecB-Halo: 355, total number of cells the RecB-Halo-Halo:290

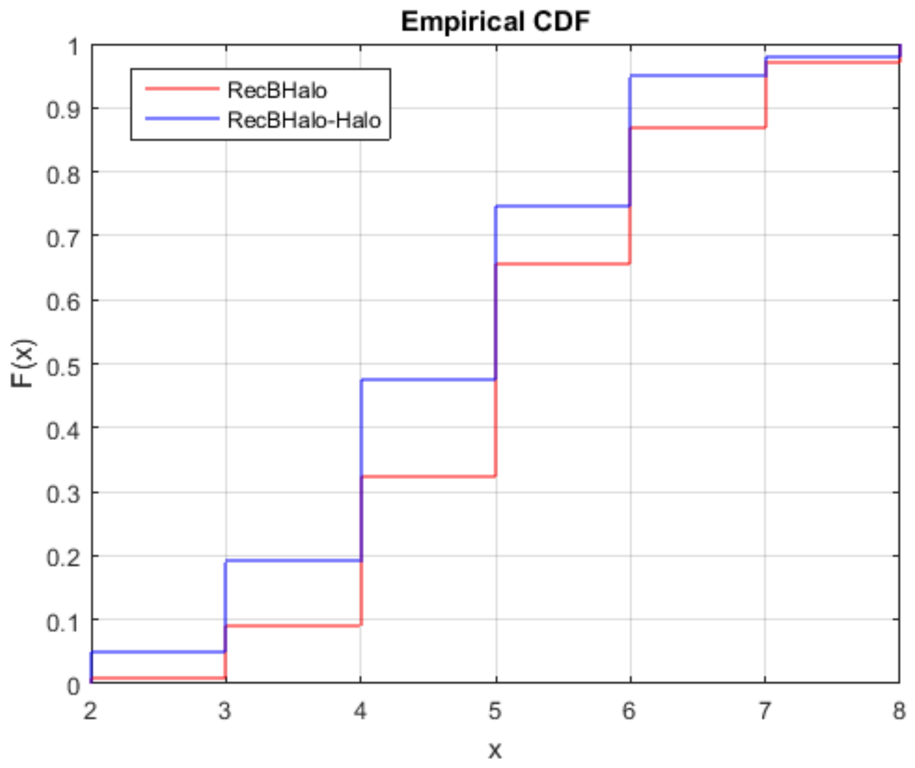


**Figure S8. RecB-Halo single tag detection is statistically equivalent to RecB-Halo-Halo double tag detection.**
Empirical cumulative distribution for RecB-Halo (red) and RecB-Halo-Halo (blue) are shown. A two-sample Kolmogorov-Smirnov test accepts the null hypothesis indicating that the data come from the same distribution (*p*-value = 0.90 > 0.05, the default level of significance).

.

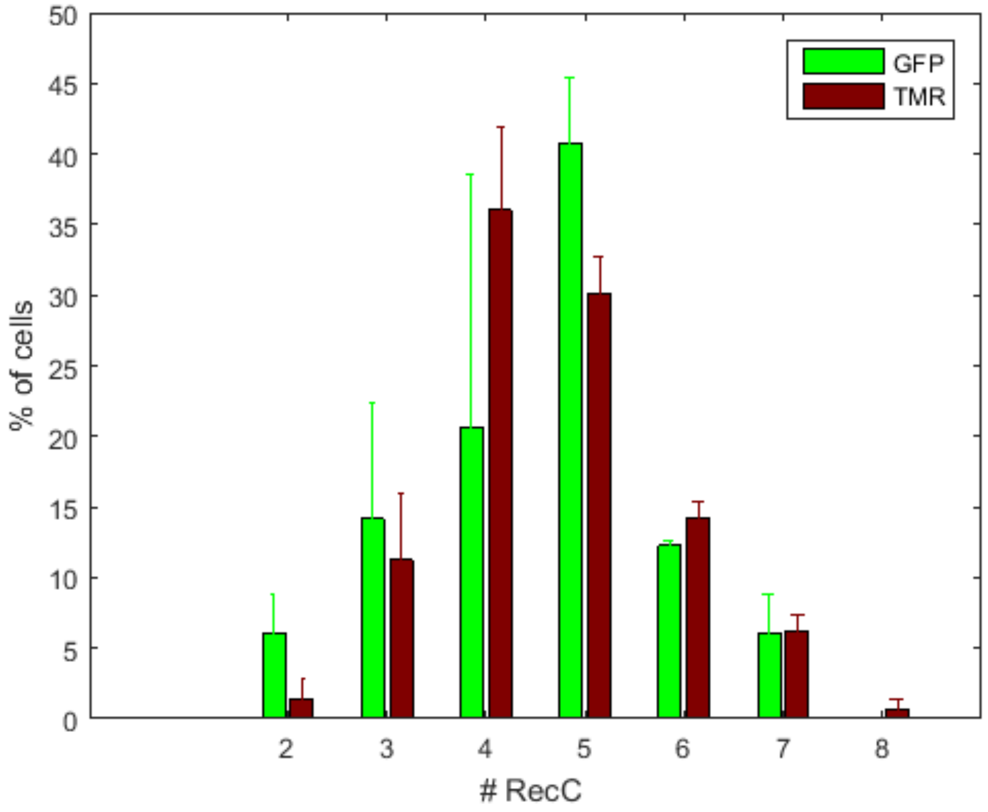


**Figure S9. RecC-Halo detection is similar to RecC-sfGFP.**
Comparison of RecC number distributions using Halo-tag based detection (red) or MACS based detection with RecC-sfGFP (green). Each distribution is the average of two experiments. Total number of cells for RecC-HaloTagLigand quantification: 584, total number of cells for RecC-GFP quantification: 50

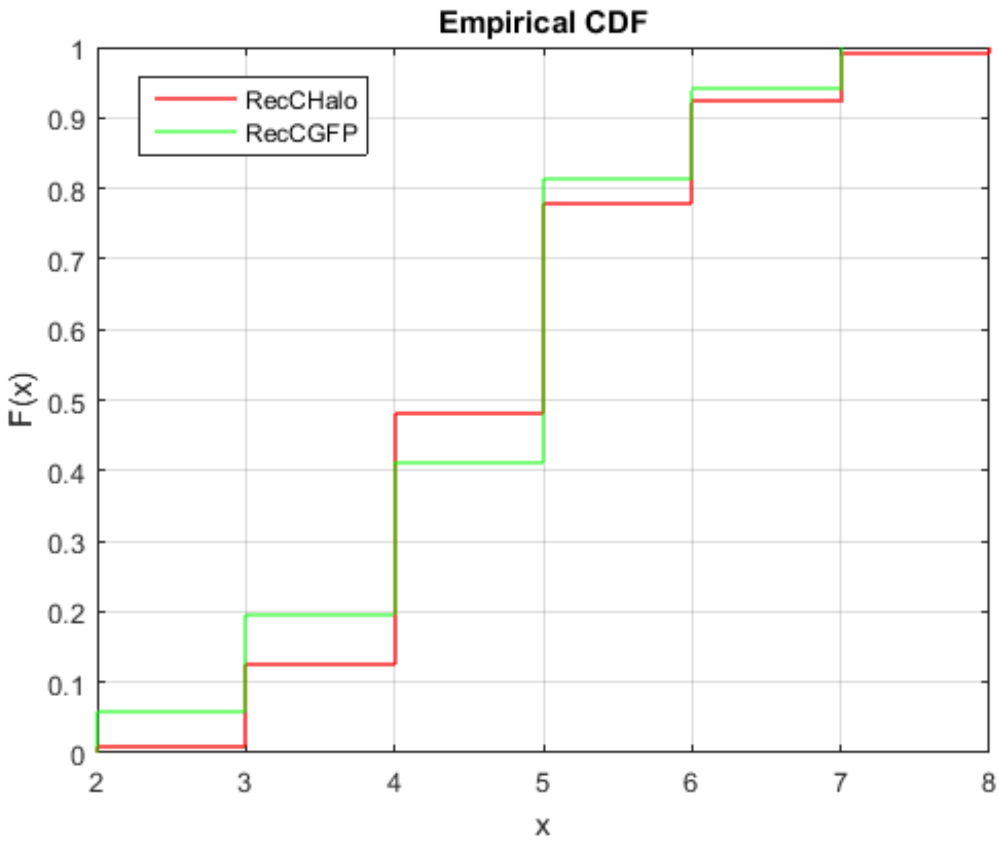


**Figure S10. RecC-Halo detection is statistically equivalent to RecC-sfGFP.** Empirical cumulative distribution for RecC-Halo (red) and RecC-sfGFP (green) are shown. A two-sample Kolmogorov-Smirnov test accepts the null hypothesis indicating that the data come from the same distribution ($p$-value = 0.94 > 0.05, the default level of significance).

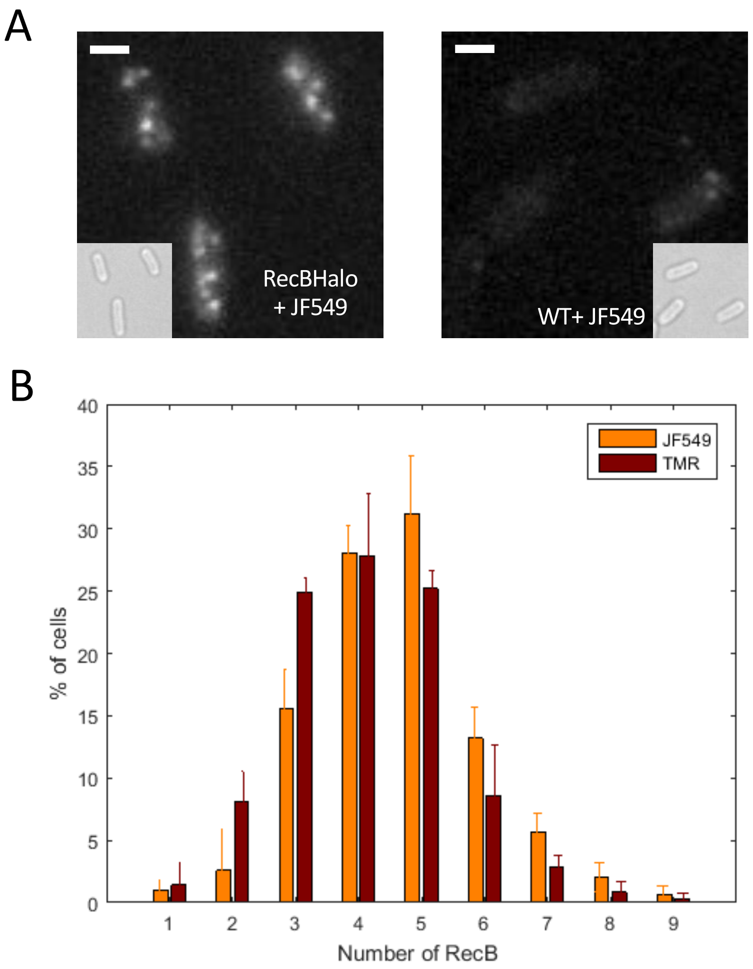


**Figure S11. RecBCD detection using JF549 is as sensitive as with HTL-TMR.** *E. coli* cells carrying the RecB-HaloTag fusion were labelled with Halo compatible JF459 dye at 5µm final concentration.

A. RecB-HaloTag appears as diffraction limited spots whilst no signal is detected in WT cells exposed to JF549, indicating that the labelling is specific. All the images are displayed with the same minimum and maximum intensity values. The maximum intensity projection of the z-stack images (from the $2^{nd}$ to the last image) is shown. Bar: 1µm

B. The distribution of RecB-HaloTag detected by HTL-TMR (dark red) or JF459 (orange) is similar indicating that both dyes have the same labelling efficiency. (Total number of cells ~ 300 for both samples)

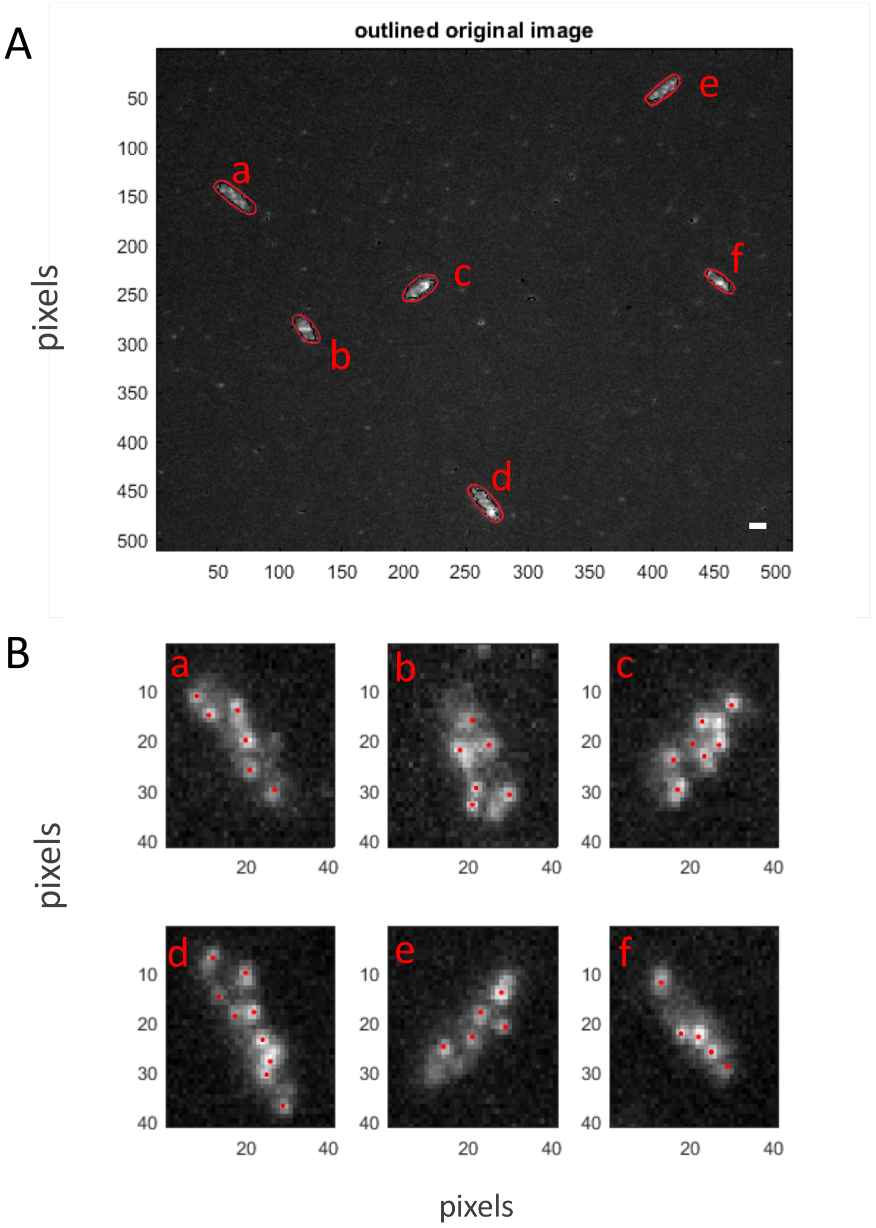


**Figure S12.** Cell segmentation and diffraction limited spots detection. A. Example image of segmented cells. The cell outline (highlighted in red as visual aid) is found by analyzing the maximum intensity projection of all the acquired z-stack. In the image, six cells were found (a to f) B. Each of the detected cells is separately analysed by defining a ROI around the cell centroid. The image is then filtered using a spatial band pass filter and diffraction limited spots are identified as local maxima within an area of 6x6 pixels.

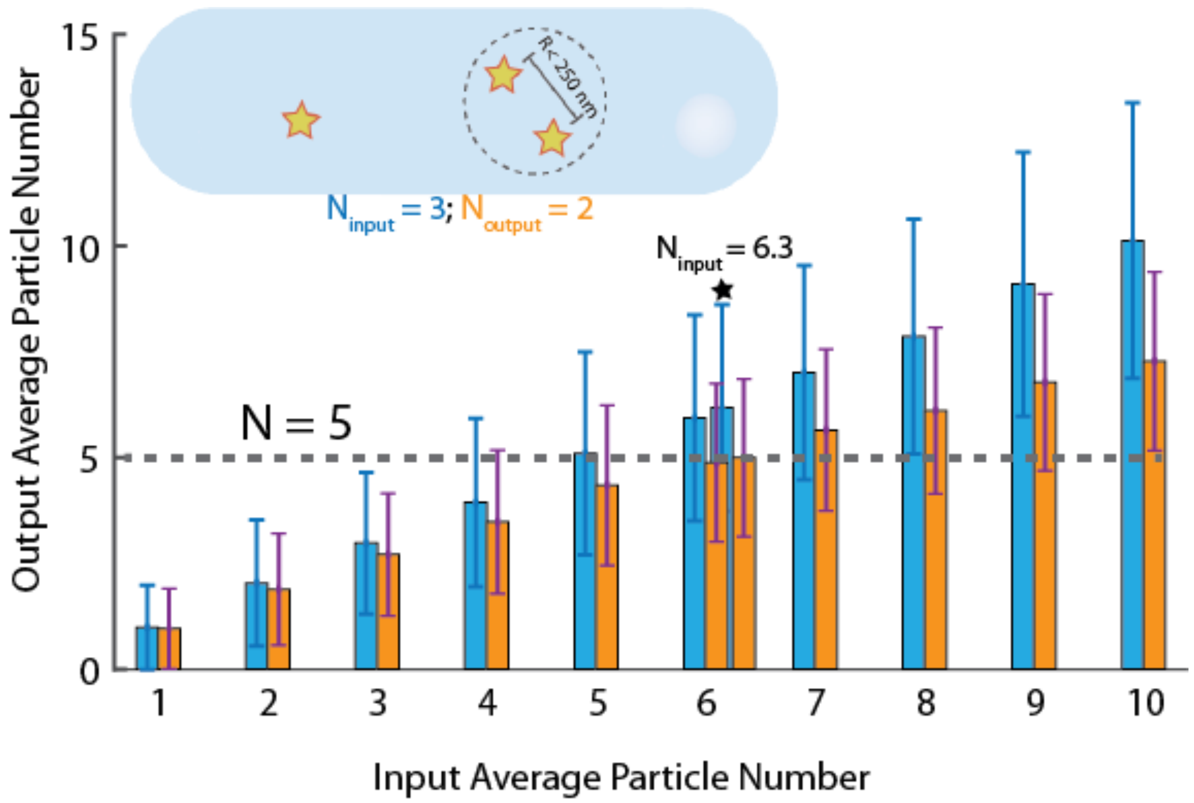


**Figure S13.** (Top right) Schematic of simulating particles in a virtual bacterial cell and undercounting from proximity and diffraction limit. Due to close proximity of two of the particles (circled), they are counted as one. This results in the detected number of particles to be 2, where the input particle number is 3. For each input average number of particles (x-axis), we have sampled 1000 numbers from a Poisson distribution with that mean, and compared the input distribution (blue; mean and standard deviation) with the detected particle distribution (orange; mean and standard distribution). For the input mean of 6.3 particles per cell, the detected mean matches the experimental average.

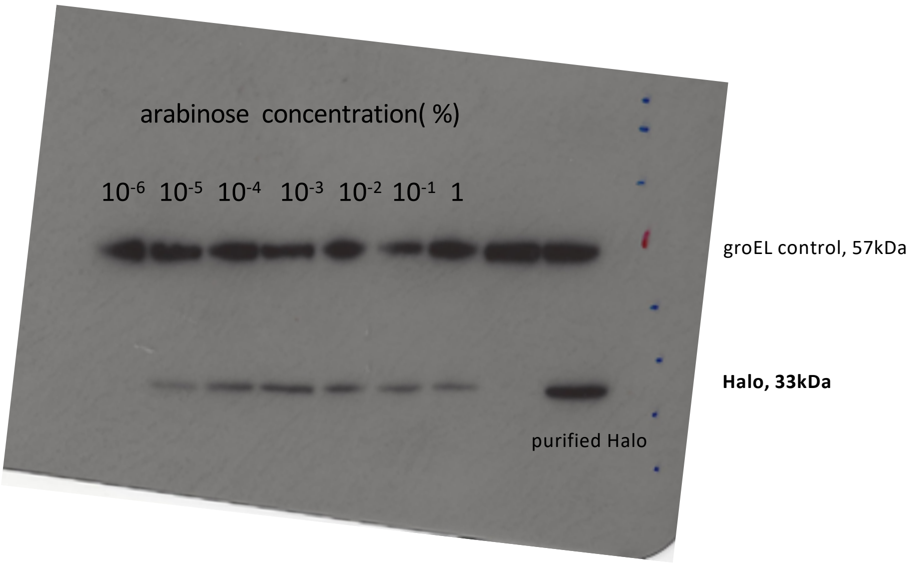


**Figure S14. Original image of the western blot shown in Fig1D.** The arabinose concentration (%) range is from $10^{-6}$ % to 1% of arabinose. In Fig 1D, the full Western blot is not shown since at $10^{-6}$%, no HaloTag expression is detected.

### Suppl. Table

### Table S1: Oligonucleotides used in this work

| Designation | Sequence (5’– 3’) |
|---|---|
| OSF1 | TTTCTCCATACCCGTTTTTTTGGGCTAGCGAATTCGAGCTAAAGAGGAGAAAGGATCCAT |
| OSF2 | CTTCTCTCATCCGCCAAAACAGCCAAGCTTGCATGCCTTAACCGGAAATCTCCAGAGTAG |
| Obh35 | GGTGCGCATCATAAAGTAAGCGGATAGATTGCGCAATTTTTATACAGCACATTCCGGGGATCCGTCGACC |
| Obh36 | TGAACAGTCGCAACGTTTCCTGTTACCGCTGTTTCGCTTTAATCAGTCAAGCGGTGGCGGTGGCAGTAA |
| Obh41 | CGCTTATGTCTATTGCTGGTCTCGGTACCCGACCTGCAAGGCGGTAATTACCCAGATGC |
| Obh46 | TGCGCCGCTACAGGGCGCGTCCCATTCGCCACCGCTCGAGATTAATATCGCGCAGCAACG |
| Obh67 | TTACTGCCACCGCCACCGCTGGAACCGCCTAGTCCAAGTAACA |
| Obh66 | CGCTCTATTTGCGCCTGTTACTTGGACTTGGACTAGGCGGTTCCAGCGGTGGCGGTGGCAGTAA |
| Obh70 | TCAGCGGGCGGGGAAAGGCGGCTCCAGGTGCTCCAGAACCGGAAATCTCCAGAGTAGAC |
| Obh69 | TCTGGAGCACCTGGAGCCGCCTTTCCCCGCCCGCTGACCGTTGA |
| Oht82 | GCCGCTTATGTCTATTGCTGGTCTCGGTACCCGACCTGCAAGGCGGTAATTACCCAGATG |
| Oht83 | CTTCCTGGCAGCCGCCTCTTTCGCTGCGGCTTCAGCCAGACCGGAAATCTCCAGAGTAG |
| Oht86 | CGAGATCGCGCGCTGGCTGTCTACTCTGGAGATTTCCGGTTCTGGAGCACCTGGAGCC |
| Oht87 | ATGCGCCGCTACAGGGCGCGTCCCATTCGCCACCGGTCGAG TTAAT TCGCGCAGCAAC |
